# Development of Broad-spectrum β-cyclodextrins-Based Nanomaterials Against Influenza Viruses

**DOI:** 10.1101/2024.03.05.583334

**Authors:** Arnaud Charles-Antoine Zwygart, Chiara Medaglia, Yong Zhu, E Bart Tarbet, Westover Jonna, Clément Fage, Didier Le Roy, Thierry Roger, Samuel Constant, Song Huang, Francesco Stellacci, Paulo Jacob Silva, Caroline Tapparel

## Abstract

In recent decades, epidemics and pandemics have multiplied throughout the world, with viruses generally being the primary agents responsible. Among these, influenza viruses play a key role, as they cause severe respiratory distress, representing a major threat to public health. To enhance the response to viral disease outbreaks, there is a need for ready-to-use broad-spectrum antivirals. We have engineered macromolecules (named CD-SA) consisting of a β-cyclodextrin (CD) scaffold modified with hydrophobic linkers in the primary face, onto which unitary sialic acid (SA) epitopes are covalently grafted, this to mimic influenza virus host receptors. In this study, we demonstrated that CD-SA, with a unitary SA, without extensive polysaccharides or specific connectivity, acts as a potent virucidal antiviral against several variants of human influenza type A and type B viruses. We also assessed the genetic barrier to resistance of CD-SA *in vitro* and successfully delayed emergence of resistance by combining CD-SA with interferon-λ1 (IFN λ1). Finally, we completed the characterization of the antiviral activity by conducting both *ex vivo* and *in vivo* studies, demonstrating a potent antiviral effect in human airway epithelia and in a mouse model of infection, higher than that of Oseltamivir, a currently approved anti-influenza antiviral.

## 1. Introduction

Over the last decades, epidemics and pandemics caused by viral pathogens have risen drastically. Globalization increases the circulation of people and goods at high volumes and speed, which offers infectious agents a constant avenue to spread worldwide [1]. It is difficult to predict which virus will trigger the next pandemic, but influenza viruses (IVs) are among the most likely candidates. IVs cause annual epidemics during the cold season and have been responsible for four pandemics in the last century [2, 3]. IV enters respiratory epithelial cells. When restricted to the upper airways, the infection causes rather mild disease, but if it spreads to the lungs it can cause viral pneumonia, with progression to acute respiratory distress syndrome and death from respiratory failure [4]. IV also disrupts the barrier functions of the respiratory epithelium, making the host more vulnerable to secondary infections by other pathogens, which significantly contribute to the morbidity and mortality of influenza [5, 6]. Given this scenario, ready-to-use antivirals active against several IV variants can relieve the human and economic burden caused by this disease.

IV is an enveloped virus, with a negative single-stranded RNA genome. The outer lipid membrane, which is taken from the host cell, is decorated with the hemagglutinin (HA) and neuraminidase (NA) proteins, which mediate viral attachment and release, respectively. IVs use HA homotrimers to recognize terminal sialic acids (SA) attached to galactose residue and thereby initiate infection [7, 8]. Human IVs bind to α2-6 linked SA, whereas avian IVs interact with α2-3 linked SA. α2-3 and α2-6 linked SA are heterogeneously distributed in the human respiratory tract. The prevalence of α2-6 linkages decreases from the upper to the lower respiratory system, until it reaches an approximately equal ratio to that of SA with α2-3 linkages in the alveoli [8, 9]. It has been reported that the characteristic structural topology rather than the nature of the linkage itself enables specific binding of HA to either α2-6 or α2-3 sialylated glycans [9, 10]. Moreover, the binding affinity of these viral proteins for sialoglycan-based receptors is enhanced by multivalent interactions [11].

In previous studies, we reported the design strategy and the mechanism of action of CD-6’SLN [12-14], an antiviral molecule that mimics the attachment moieties of HA from human-adapted viruses. This compound is made of a β-cyclodextrin core (CD) grafted with long alkyl chains terminating in 6’SLN (6’-Sialyl-N-acetyllactosamine, a trisaccharide terminating with an α2-6 SA) (Fig.1A). Such an approach allowed trapping and irreversibly damaging viruses, thereby preventing them from attaching to host cells. This resulted in potent antiviral activity in immortalized cell lines, in *ex vivo* reconstituted human respiratory epithelia and in mice [13, 15].

**Figure 1.**
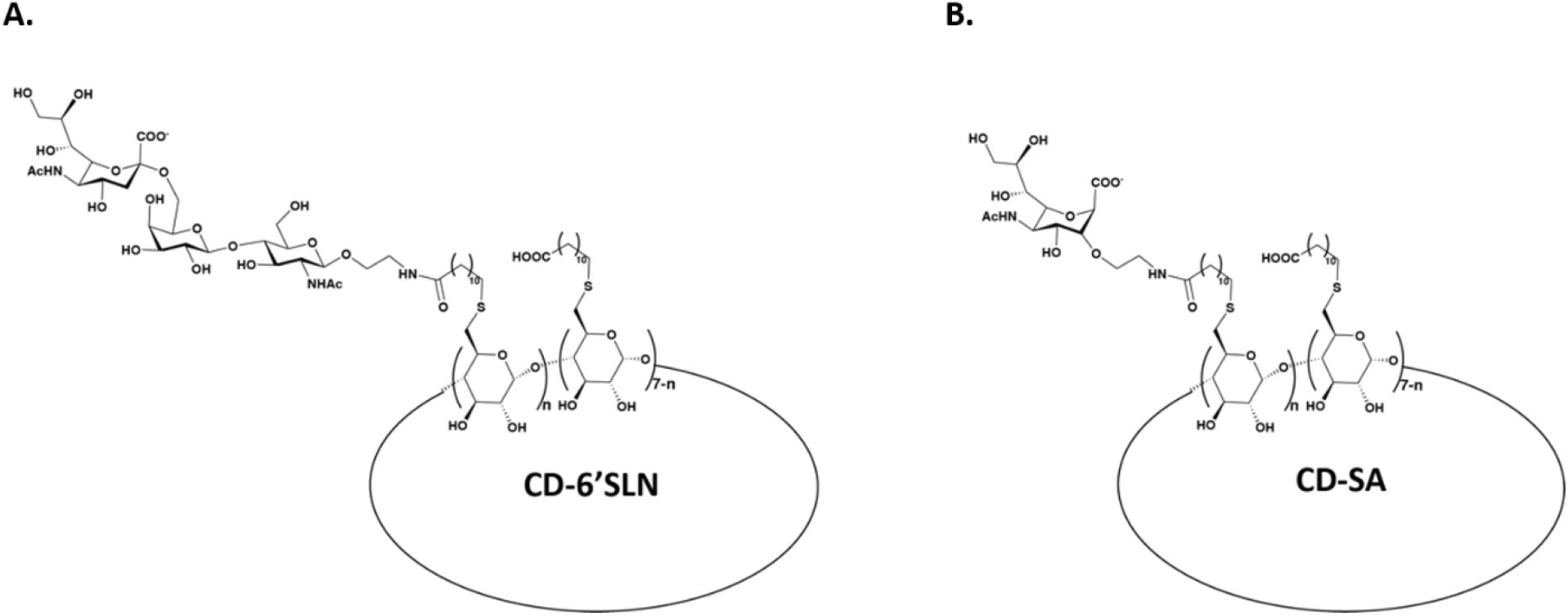
Schematic representation of CD-6’SLN and CD-SA. A β-CD core is modified on its primary face with hydrophobic undecyl linkers exposing either 6’SLN (6’-Sialyl-N-acetyllactosamine), a trisaccharide terminating with an α2-6 sialic acid for CD-6’SLN (**A**) or a single sialic acid residue for CD-SA (**B**) [12].

In the current study we investigated the efficacy and the mechanism of action of CD-SA, a novel CD based molecule, grafted with long alkyl chains terminating in a single residue of SA (Fig.1B). In contrast to CD-6’SLN, the molecular arrangement of CD-SA should allow binding to the HA pocket of a broader range of IV strains. Additionally, it may present a higher barrier to antiviral resistance by preventing the receptor switch from α2-6 to α2-3 SA. We tested and confirmed the antiviral activity of CD-SA against several IV strains and demonstrated that its genetic barrier to antiviral resistance is indeed higher than that of CD-6’SLN. Lastly, we proved CD-SA antiviral activity both *ex vivo* and *in vivo*.

## 2. Materials and methods

The synthesis of CD-SA as well as more detailed methodologies are available in the supplementary file.

### 2.1 Cells and tissues

Madin-Darby canine kidney (MDCK, ATCC) were cultured in Dulbecco’s Modified Eagle Medium (DMEM) supplemented with Gluta-MAX™, Sodium Pyruvate, Phenol Red, 10% FBS and 1% P/S and grown at 37°C, with 5% CO2. Calu-3 cells were cultured in Minimum Essential Medium (MEM) supplemented with Gluta-MAX™, 10% FBS, Phenol Red, 1% Hepes, 1% Non-Essential Amino Acids, 1% penicillin/streptomycin, and 1% Sodium pyruvate and grown at 37^0^C in an atmosphere of 5% CO2.

Human airway epithelia (HAE) (MucilAir™, Epithelix, Plan-les-Ouates, Switzerland) were reconstituted from freshly isolated primary human nasal polyp epithelium collected either from 14 different donors, upon surgical nasal polypectomy, or from individual donors. The donor patients presented no atopy, asthma, allergy or other known comorbidities. The study was conducted according to the Declaration of Helsinki on biomedical research (Hong Kong amendment, 1989), and the research protocol was approved by the local ethics committee. The tissues were maintained at the air-liquid interface according to the manufacturer’s instructions [5].

### 2.2 Viruses

Human A/Netherlands/602/2009(H1N1) influenza virus (H1N1/pdm09) and human B/Wisconsin/01/2010, Yamagata lineage (Influenza B virus) were provided by Prof. Mirco Schmolke (University of Geneva, Geneva, Switzerland). Viral stocks were amplified and titrated in MDCK cells by plaque assay. Briefly, cells infected with serially diluted viruses were overlaid with MEM containing 0.3% BSA, 0.9% bacto agar, and 1μg/ml N-tosyl-L-phenylalanine chloromethyl ketone (TPCK)-treated trypsin. After 48 h at 37°C, cells were fixed with 4% formaldehyde and stained with 0.1% crystal violet, to calculate the number of plaque forming units (PFU). Plaque assays were run in biological duplicates.

For *in* vitro and ex *vivo* experiments, Human A/Switzerland/2022(H1N1) and human A/Switzerland/2022(H3N2) were provided by the diagnostic laboratory from the HUG (Hôpitaux Universitaires de Genève) and viral stocks were prepared in MucilAir tissues as previously described [14].

For *in vivo* experiments, mouse adapted influenza A/California/04/2009(H1N1pdm) was kindly provided by Dr. Elena Govorkova (St. Jude Children’s Research Hospital, Memphis, TN). The virus stock was prepared and tittered by cell culture infectious dose 50% assay (CCID50) in MDCK cells.

### 2.3 Cell viability assay

Cell viability was assessed by MTT assay as described in [16] while viability in *ex vivo* tissues was assessed by Resazurin treatment as described in [17].

### 2.4 Antiviral dose response assays and time of addition assays

The viral inhibition assay was performed as previously described [12]. A dose range of molecules was pre-incubated with the virus and the number of infected cells was calculated by immunocytochemistry (ICC). The effective concentration (EC) 50 were calculated by nonlinear regression analysis [log(inhibitor) vs. response – Variable slope (four parameters)] in GraphPad prism Software. For the cell pretreatment, cells were initially exposed to CD-SA for 1 h, followed by washing and virus inoculation. In the cotreatment method, CD-SA was simultaneously added with the virus to the cells. For the post-infection condition, CD-SA was added to the cells 1 h post infection (hpi). Number of infected cells was calculated by ICC 16 hpi. All results are presented in GraphPad Prism (GraphPad Software version 8.0, San Diego, CA, USA) as mean from two independent experiments performed in duplicate.

### 2.5 Ex vivo antiviral testing assay

MucilAir tissues were infected apically with 1E5 RNA copies/tissue of H1N1/pdm09 in MucilAir medium (Epithelix), at 37°C for 5 h. After inoculum removal, the apical side was washed three times with PBS^++^ and 30µg of CD-SA11 or CD-6’SLN were administered apically in 30µl of MucilAir medium. Viruses released apically were harvested daily and 30µg of fresh drug was added. Viral loads were assessed by RNA extraction and RT-qPCR.

### 2.6 Virucidal assay

1E6 PFU of each variant was incubated with one EC99 of CD-SA for 3 h at 37°C. The mixture was then serially diluted in serum-free DMEM containing 1% penicillin/streptomycin and transferred on MDCK cells, seeded in a 96-multiwell plate. After 1 h, the mixture was removed and fresh medium was added. At 16 hpi viral titers were evaluated by ICC, and percentages of infection were calculated compared to the untreated control (incubated 3 h at 37°C with buffer).

### 2.7 RNA exposure assay

H1N1/pdm09 (1E6 PFU/ml) was incubated with 100μg/ml of the molecules in DMEM medium containing 1% penicillin/streptomycin at 37°C for 3 h, and half of the mixture was treated with RNase, while the other half with buffer. Briefly, 1.1μl of 10mg/ml RNAse A (EN0531, Thermo Fisher Scientific, Waltham, MA) or 1.1μl of MilliQ water was added into 10μl of PBS buffer and mixed with 100μl of a diluted (1:20) viral-molecule solution. After 30min at 37°C, 500μl of GTC lysis buffer (Omega bio-tek, Norcross, GA) was added, and RNA was extracted and quantified by RT-qPCR.

### 2.8 TEM assay

Transmission Electron Microscopy (TEM) was performed at the Dubochet Center for Imaging (DCI Geneva). Concentrated H1N1/pdm09 stock (1E7 PFU) was incubated with control medium or 1.7µg/ml of non-functionalized (non-modified β-cyclodextrins) CD or CD-SA for 1 h at 37^°^C. Following fixation with glutaraldehyde 4% for 24 h at 4^°^C, viruses were negatively stained with uranyl acetate on carbon-coated copper grids and analyzed by TEM at 29’000 times magnification with a TEM microscope at 120kV (Tecnai G2, FEI, Eindhoven).

### 2.9 Preparation of R18-labeled H1N1/pdm09 and fluorescence release assay

2.33μl (3.12μmol/ml) of ethanolic R18 (octadecyl rhodamine B chloride, Sigma Adrich) was added to a 255μl H1N1/pdm09 suspension [2mg/ml of viral protein quantified using the modified Lowry protein assay (Thermo Scientific™ Pierce™ Modified Lowry Protein Assay Kit, Thermo Fisher Scientific, Waltham, MA)]. The mixture was incubated in the dark for 1 h at RT. Non-incorporated fluorophores were removed using Zeba™ Spin Desalting columns (7K MWCO, Thermo Fisher Scientific, Waltham, MA) with 10mM TES, 150mM NaCl (pH 7.4) as elution buffer. The protein concentration of the R18-labeled virus was re-assessed using the modified Lowry assay. Two μg/ml of R18-labeled H1N1 in 1% penicillin/streptomycin DMEM medium were incubated with 100μg/ml of non-functionalized CD, 1% Triton X-100 or 100μg/ml CD-SA and incubated for 3 h. Fluorescence was measured using a Tecan Infinite 200 plate reader with excitation at 560nm and emission at 590nm.

### 2.10 Antiviral resistance assay

H1N1/pdm09 was passaged 9 times in Calu-3 cells in the presence of increasing concentrations of CD-SA, CD-6’SLN, and a combined therapy of CD-SA with IFNλ1 or in presence of serum-free MEM only. Viral loads were quantified by titration in MDCK cells. Viruses presenting increased viral loads were sequenced and the drug sensitivity was checked via a dose-response assay (§2.4).

### 2.11 RT-qPCR analysis, viral RNA quantification and sequencing

Viral RNA was quantified using RT-qPCR as previously described [14]. In the RNA exposure assay, extracted RNAs were reverse transcribed, purified and sequences were obtained through Fasteris DNA sequencing service (Geneva) and analyzed with Geneious software.

### 2.12 In vivo mouse model

Female 18-20g BALB/c mice were obtained from Charles River Laboratories (Wilmington, MA). Mice (n=5 per group for toxicity assay or 10 per group for efficacy assay) were anesthetized by intra-peritoneal (i.p.) injection of ketamine/xylazine (50/5mg/kg). For toxicity assays mice were administered intranasally (i.n.) with 50μl of daily CD-SA treatment (up to 80mg/kg) for 6 days. For efficacy assays, anesthetized mice were inoculated by the i.n. influenza challenge (30 CCID50 in 75μl of PBS). Daily treatments of 40 or 7.5mg/kg of CD-SA or the vehicle placebo sterile saline were administered beginning 1 day and 2 days pi. Oseltamivir (OS) phosphate (30mg/kg/day) prepared in sterile water for oral gavage (per os) was administered in 100μl for 5 days starting 2 h before infection as the positive control. Individual weights were recorded every day beginning on the day of virus challenge and mice were observed daily for survival. Survival curves were compared by the Mantel-Cox log rank test. Mean day of death (MDD) comparisons were made by one-way ANOVA with Dunnett’s multiple comparisons test. Differences in the number of survivors between compound-treated and placebo groups were analyzed by the Fisher’s exact (two-tailed) test. Calculations were made using Prism 9 (GraphPad Software, San Diego, CA). This study was conducted in accordance with the approval of the Institutional Animal Care and Use Committee of Utah State University and done in the AAALAC-accredited Laboratory Animal Research Center of Utah State University.

## 3. Results

### 3.1. CD-SA inhibits A/Netherlands/602/2009(H1N1) with a virucidal mechanism of action

The antiviral effect of CD-SA (Fig.1A-B) against IV A/Netherlands/602/2009(H1N1) (named hereafter H1N1/pdm09) was tested by pretreating the virus 1 h with the molecule at 37°C before infection of MDCK cells. In these conditions, the EC50 of CD-SA was of 4.75ng/ml (Fig.2A) which is ≈40 times lower than that of CD-6’SLN (180ng/ml) against the same virus [12]. Of note, both compounds have a cytotoxic concentration 50 (CC50) >200μg/ml (Table1). To assess the antiviral mechanism of action, a dose range of CD-SA (from 0.02μg/ml to 100μg/ml) was administered at different times relative to the infection. The compound was either applied directly on cells before infection (cell pretreatment condition), was administered at the time of infection (cotreatment condition), or was added 1 h after infection (post-treatment condition). As shown in Fig. 2B, none of these conditions was as effective as the virus pretreatment condition (Fig. 2A). The cotreatment was the second most effective condition (96.55 ± 15 % inhibition at 100μg/ml), while a weak inhibition was observed in both the cell pretreatment (30 ± 19.5 % inhibition at 100μg/ml) and the post-treatment conditions (19,5 ± 16.9 % inhibition at 100μg/ml). These results indicate that the antiviral effect of CD-SA is based on the compound interaction with the virus, rather than a cell-mediated effect.

**Table 1.** Dose response and virucidal activity of CD-SA and CD-6’SLN in MDCK cells against several IVA and IVB strains. The cytotoxic concentration 50 (CC50) and effective concentrations 50 (EC50) were calculated with MTT assay and dose response assay by nonlinear regression analysis in GraphPad prism. The presence or absence of virucidal activity was assessed via a virucidal assay. Results present the mean of two independent experiments performed in duplicate [95% Confidence interval (CI)]. 1μg/ml of CD-SA is equivalent to 290nM.

| VIRAL STRAIN | CD-SA<br>AND CD-6'SLN | CD-SA |  |  | CD-6'SLN |  |  |
| --- | --- | --- | --- | --- | --- | --- | --- |
|  | CC50<br>[µg/ml] | EC50<br>[µg/ml] [CI] | SI | Virucidal<br>Activity | EC50<br>[µg/ml] [CI] | SI | Virucidal<br>Activity |
| A/Netherlands/602/2009(H1N1) | >200 | 0.0048<br>[0.0036 to 0.0062] | >41666 | + | 0.12<br>[0.11 to 0.14] | >1666 | + |
| A/Switzerland/2022(H1N1) | >200 | 7.14<br>[5.37 to 10.31] | >26.8 | + | 5.96<br>[4.54 to 8.3] | >30.48 | + |
| A/Switzerland/2022(H3N2) | >200 | 10.54<br>[6.85 to 15.24] | >19 | - | 7.54<br>[5.3 to 10] | >26.52 | - |
| B/Wisconsin/2010 (Yamagata) | >200 | 27.8<br>[15.92 to 51.79] | >7.2 | + | 4.76<br>[3.1 to 6.4] | >42 | + |

**Figure 2.**
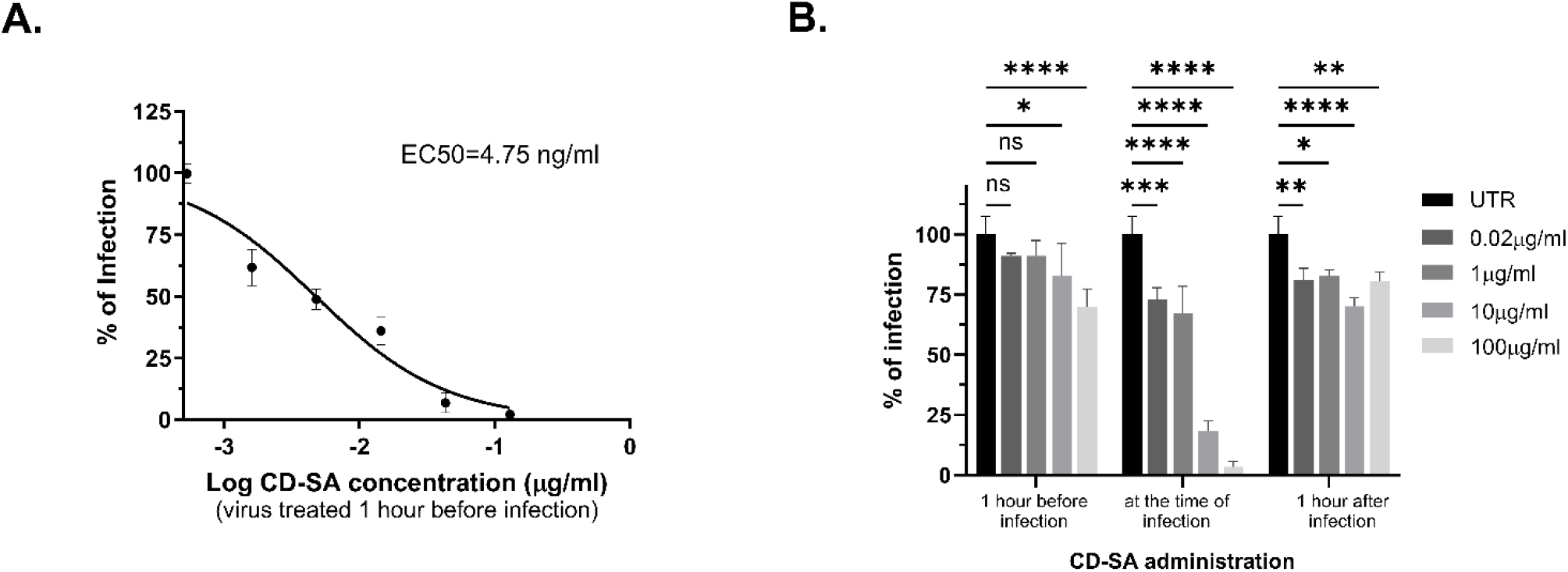
Antiviral mechanism of CD-SA. Dose-response assay of CD-SA against H1N1/pdm09 in virus pretreatment condition (**A**) and efficacy depending on time of administration (**B**). In **A**, CD-SA and viruses were incubated for 1 h at 37°C before infection. In **B**, MDCK cells were either pretreated with CD-SA for 1 h before infection; infected with virus and treated with CD-SA at the same time; of finally, treated with CD-SA 1 h after viral inoculation. Percentages of infection are shown as mean and SD. P-values (*, ≤ 0.1; **, ≤ 0.01; ***, ≤ 0.001; ***, ≤ 0.0001) were calculated using two-way ANOVA.

We then assessed whether CD-SA, like other CD-based antivirals bearing long aliphatic linkers [12, 18], was virucidal (Fig.3). We first performed a virucidal assay, incubating concentrated H1N1/pdm09 with CD-SA (1.7μg/ml) for 3 h [12]. The virus-drug complex was then serially diluted before infection and the viral titers were determined by ICC. As shown in Fig.3A, the viral infectivity was greatly decreased even at dilutions leading to a final drug concentration lower than the EC50, confirming an irreversible viral inactivation. We then checked whether binding to CD-SA would induce virus disruption and consequently viral genome release, using an RNA exposure assay [13]. Briefly, CD-SA (100μg/ml) or the surfactant sodium dodecyl sulfate (SDS) were incubated with H1N1/pdm09 for 3 h at 37°C and treated with RNase or buffer. After lysis and RNA extraction, intact genomes were quantified by RT-qPCR and the cycle threshold difference (ΔCt) between RNAse treated and untreated samples were calculated. We found that CD-SA did not significantly change the ΔCt compared to the untreated virus, while the positive SDS control resulted in an approximately 8-fold increase of the ΔCt (Fig.3B). These results show that unlike SDS, CD-SA does not make the RNA accessible to RNase activity. However, this does not mean that the viral envelope is intact, as the viral nucleoprotein provides another layer of protection for the viral RNA.

To study the impact of CD-SA on the viral envelope, we imaged its effect on H1N1/pdm09 structure by transmission electron microscopy (TEM) (Fig.3C). H1N1/pdm09 was incubated for 1 h at 37°C with 1.7μg/ml of non-functionalized CD, or the virucidal CD-SA, and negatively stained with uranyl acetate. As shown on the TEM images, the viral envelope is destroyed by the incubation confirming the virucidal effect of CD-SA (Fig.3C). These results were also supported by a fluorescent release assay (Fig.3D), in which the envelope of H1N1/pdm09 was labelled with the amphiphilic octadecyl rhodamine B chloride (R18) dye. The dye fluorescence is quenched at high concentrations on the membrane and released at low concentrations upon membrane disruption. R18-labeled viruses were incubated with CD-SA, non-functionalized CD and Triton X-100 (known to disrupt influenza membrane) for 3 h and the relative fluorescence unit (RFU) was measured. As shown in Fig.3D, both the virucidal CD-SA and the positive Triton X-100 control significantly increased the RFU compared to the untreated virus, while the non-functionalized CD showed no difference.

**Figure 3.**
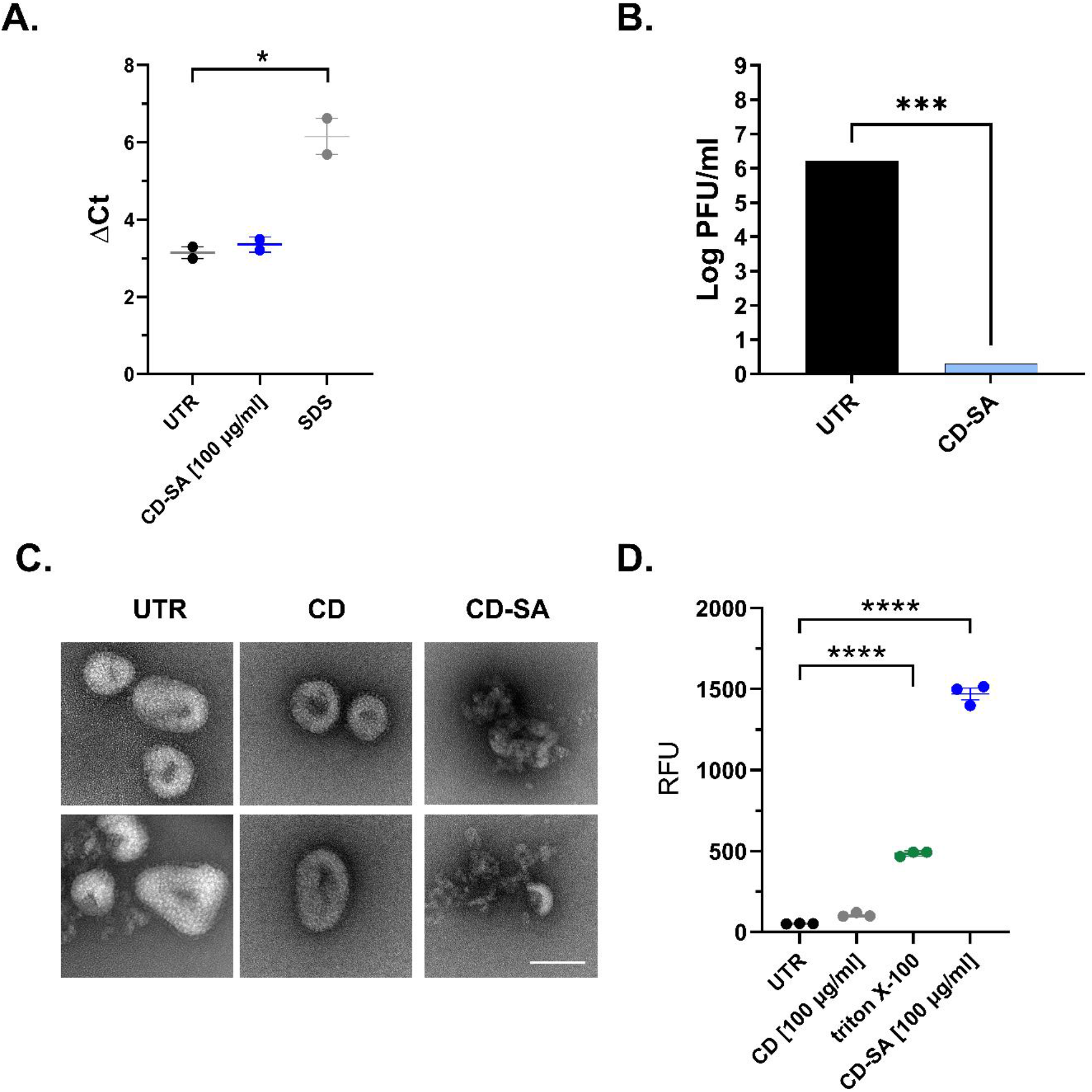
Virucidal activity of CD-SA against H1N1/pdm09 A. RNA exposure assay: Cycle threshold difference in H1N1/pdm09 RT-qPCR between RNase-treated virus molecule mixture and buffer-treated virus molecule mixture. The p-value (*, ≤ 0.05) was determined using a two-tailed unpaired t-test. **B**. Virucidal assay: H1N1/pdm09 and CD-SA or the CD-SA vehicle (Untreated condition = UTR) were incubated for 3 h at 37°C and serially diluted onto MDCK cells and viral titers were evaluated by ICC. A reduction of viral titer by >2 log indicates a virucidal mechanism of action [12]. The p-value (***, ≤ 0.0001) was determined using an unpaired t-test. **C**. Negative staining electron microscopy: H1N1/pdm09 was incubated with control medium or 1.7μg/ml of non-functionalized CD, or virucidal CD-SA for 1 h at 37°C and processed for TEM. Scale bar is equivalent to 50 nm. **D**. Envelope integrity assay: relative fluorescence unit of R18-labeled H1N1/pdm09 measured after 3 h incubation with non-functionalized CD, triton X-100 and CD-SA. P-values (****, ≤ 0.0001) were determined using one-way ANOVA.

### 3.2. CD-SA displays direct antiviral activity against different influenza A and B strains

As CD-SA exposes a unique SA epitope covalently bound to an undecyl linker, it should bind the HA pocket of several IV strains. Hence, we compared the spectrum of antiviral activity of CD-SA and CD-6’SLN that share the same epitope presentation. We tested these molecules in a dose-dependent manner and compared their virucidal activity against a range of influenza variants from groups A and B. As shown in Table1, the EC50 and virucidal activity of the two compounds are comparable against the different strains tested although the EC50 of CD-SA is lower against H1N1/pdm09 (0.0048µg/ml versus 0.12µg/ml) but higher against IVB (27.8µg/ml versus 4.76µg/ml). Surprisingly none of the compounds show a virucidal activity against a recent A/Switzerland/2022(H3N2) strain, while they showed virucidal activity against A/Singapore/37/2004 (H3N2) ([12]; Data not shown).

### 3.3. CD-SA displays antiviral activity in a relevant respiratory tissue culture model

In our previous work, we showed that CD-6’SLN blocks IV infection in *ex vivo* reconstituted human respiratory tissues [15]. Here, we conducted the same test for CD-SA, with CD-6’SLN serving as control. We infected reconstituted human airway epithelia (HAE) with 1E5 RNA copies of H1N1/pdm09 and treated them daily for 5 days, starting at 1 day post-infection. We then monitored, every day, the amount of virus released from the apical tissue side (Fig.4A). We observed a significant reduction of viral replication in treated versus untreated tissues, confirming the therapeutic potential of CD-SA in this relevant tissue culture model (Fig.4B). Of note, no change in metabolic activity was observed in treated versus untreated tissues at this compound concentration (Fig. 4B, dashed lines).

**Figure 4.**
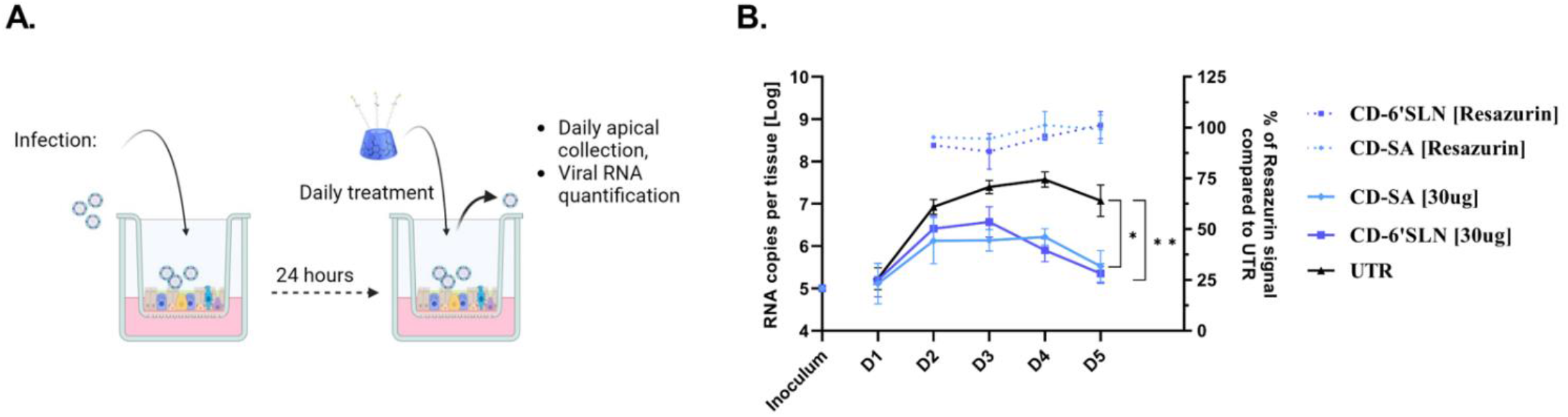
Antiviral efficacy of CD-SA and CD-6’SLN against H1N1/pdm09 in *ex vivo* human airway epithelia (HAE) reconstituted at the air-liquid interface. **A**. Schematic of *ex vivo* HAE infection and treatment. **B**. Tissues were infected apically and antivirals were administered from 1 dpi to 4 dpi. Cell viability of uninfected treated controls was assessed by resazurin assay. Viral loads, expressed in RNA copies/tissue, were quantified by RT-qPCR from daily collected apical washes. The results represent the mean and SD from two independent experiments. Statistics were obtained by analyzing the AUC (Area Under the Curve). *, p ≤ 0.1; **, p ≤ 0.01

### 3.4. CD-SA genetic barrier to resistance *in vitro*

We next compared the genetic barrier to resistance to CD-SA or CD-6’SLN during 9 serial passages of H1N1/pdm09 in Calu-3 cells (Sup Fig.1). Calu-3 were used as they express proteases which allows multiple cycles of infection without addition of trypsin [19]. One μg/ml of compound was administered 1 hpi and the drug concentration was doubled at each passage (up to 173µg/ml). The starting concentration of 1µg/ml was chosen as it still enables good viral replication despite antiviral pressure, thus allowing emergence of antiviral resistance. In addition to these compounds and as IFNλ1 was shown to delay the emergence of resistance to oseltamivir (OS) [16], we also tested CD-SA combined with IFNλ1 in this assay. In this case, the cells were pre-treated with IFNλ1 (from 15ng/ml to 960ng/ml) 24 h before infection and then again at 1 hpi together with CD-SA as previously performed [16]. An untreated control was added to monitor cell adaptation which could spontaneously occur during the passages. At 48 hpi, the infectious viral titers were measured by plaque assay in MDCK cells (Sup Fig.1).

An increase in viral titer was observed already at passage 2 (P2) under CD-6’SLN treatment (Sup Fig.1B). In contrast, a comparable and significant loss of effectiveness was observed only at P5 for CD-SA (Sup Fig.1, Table2), indicating that CD-SA presents a higher genetic barrier to antiviral resistance. Finally, co-treatment with IFNλ1 prevented emergence of resistance against CD-SA over all 9 passages.

**Table 2.** Emergence of resistance of H1N1/pdm09 after passages in presence of increasing concentrations of CD-6’SLN, CD-SA or a combined CD-SA/IFNλ treatment.

| Drug selection | Passage<sup>a</sup> | HA mutation<sup>b</sup> | CD-6'SLN EC<sub>50</sub> [μg/ml]<sup>c</sup> | CD-SA EC<sub>50</sub> [μg/ml]<sup>c</sup> |
| --- | --- | --- | --- | --- |
| Untreated | 4 | D127E | 2.7 | 2.61 |
| CD-6'SLN | 2 | Q223Q/R |  | - |
|  | 3 | Q223R | > 100 | - |
| CD-SA | 4 | D127E | - | 13.47 |
|  | 6 | K211N/R221K | - | 128.3 |
|  | 8 | K211N/R221K | - | 214.5 |
| IFNλ1 + CD-SA | 4 | D127E | - | 3.4 |
|  | 8 | K154E | - | 3.96 |

<sup>a</sup>Passage number where mutations were detected. <sup>b</sup>Mutations numbered according to H1 pdm09 numbering [20].
<sup>c</sup>Results obtained from duplicate experiments.

Next, we sequenced the HA gene of variants presenting diminished sensitivity to the compounds as well as the control virus. We identified a mutation present in the HA of all variants from passage 4 (D127E) which is very likely due to an adaptation to Calu-3 cells. Resistance to CD-6’SLN was associated with the HA Q223R mutation which became dominant at P3, already identified in our previous study [16] and known to change receptor preference from human α2-6 to avian α2-3 linked SA [21]. Of note, H1N1/pdm09 is mice-adapted and therefore has a higher affinity for α2-3 bound SA that predominates in the mouse respiratory tract. A significant increase in the EC50 of CD-SA was associated with the K211N/R221K mutations detected at P6. The role of these two mutations is less well established. R221K, along with five other mutations, has been shown to increase the virulence of pandemic H1N1 in mice, a phenotype associated with enhanced binding to α2-3 linked SA and decreased binding to α2-6 linked SA [22]. Finally, the K154E mutation identified at P6 in the combined treatment did not cause a significant shift in the EC50 and probably emerged randomly.

### 3.5. Antiviral activity of CD-SA in mice

We then investigated CD-SA effect *in vivo*. Toxicity experiments were performed with 5 to 80mg/kg of CD-SA administered intranasally (i.n.) and for 6 days. Up to 10% weight loss was observed between day 5 and 7 (Sup. Fig 2) with the 60 and 80mg doses but mice weight returned to normal afterwards. For efficacy assay, female BALB/c mice (10 per group) were infected i.n. with 30 CCID50 of A/California/04/2009 and then treated once daily i.n. and for 5 days, with 7.5mg/kg or 40mg/kg of CD-SA starting from 1 or 2 dpi (Fig.5). A group of mice treated per os twice daily for 5 days with 30 mg/kg/day of OS (starting 2 h before infection) was added as a positive control. Contrary to OS treatment that only delayed mouse death by 2 days (no survival observed), CD-SA significantly increased survival rate when administered starting from both 1 dpi (7.5mg/kg, 40% of survival; 40mg/kg, 60% of survival; p < 0,001) or 2 dpi (7.5mg/kg, 70% of survival; 40mg/kg, 70% of survival; p < 0,001). Interestingly, survival rate was higher for mice treated after 2 days and the same survival rate was observed for mice were treated with 7.5 or 40mg/kg of CD-SA starting at 2 dpi.

## 4. Discussion

Influenza presents an important pandemic risk and IV epidemics cause severe disease and death worldwide. The center for disease control (CDC) estimates that, each year, influenza is responsible for 100’000 to 710’000 hospitalizations and 5 to 50’000 deaths in the USA [23]. Therapies are still wanting, with seasonal vaccines at the forefront and limited antiviral treatment alternatives. To tackle these challenges, novel anti-IV antivirals targeting HA, the most abundant IV surface protein, have been proposed. HA forms a weak bond with oligosaccharides terminating with SA (N-acetylneuraminic acid, Neu5Ac) [24], which prompted researchers to opt for multivalent macromolecular strategies. This has led to several macromolecules that present SA such as polymer scaffolds [25, 26], DNA-based macromolecules [27], virus-like particles [28], micelles and lipid bilayers [29-31], gold nanoparticles [32], and protein scaffolds [33-35], among other sialylated macromolecular decoys for IVs. Unfortunately, *in vivo* efficacy has not been demonstrated or addressed for most of these molecules. One possible explanation is that these glycloclusters represent a virustatic decoy for IV that does not destroy the infective particle. Hence, as the concentration of the drug decreases, infective viruses are free to infect host cells. Unlike all these approaches, CD-SA is based on a molecular architecture that exerts a hydrophobic force on the virus, leading to irreversible damage, rendering it non-infective [12, 18, 36]. This property sets CD-SA aside from most other glycoclusters and sialylated molecules.

Here, we show a strong antiviral efficacy against H1N1 pandemic strains *in vitro, ex vivo* and *in vivo*. Our *in vivo* results demonstrate the efficacy of CD-SA in rescuing mice from a lethal IV infection when added 24 or 48 h post-infection, while OS proves ineffective even if given before infection and twice daily afterwards (Fig.5). The therapeutic window of CD-SA is thus longer than that of a shedding inhibitor, likely due to the virucidal action directly on infective viral particles. The *in vivo* results also show that the dose range and time of administration of CD-SA can be optimized as the lowest dose of 7.5mg/kg/day, when administered at 2 dpi, rescued as many animals as the dose of 40mg/kg/day.

**Figure 5.**
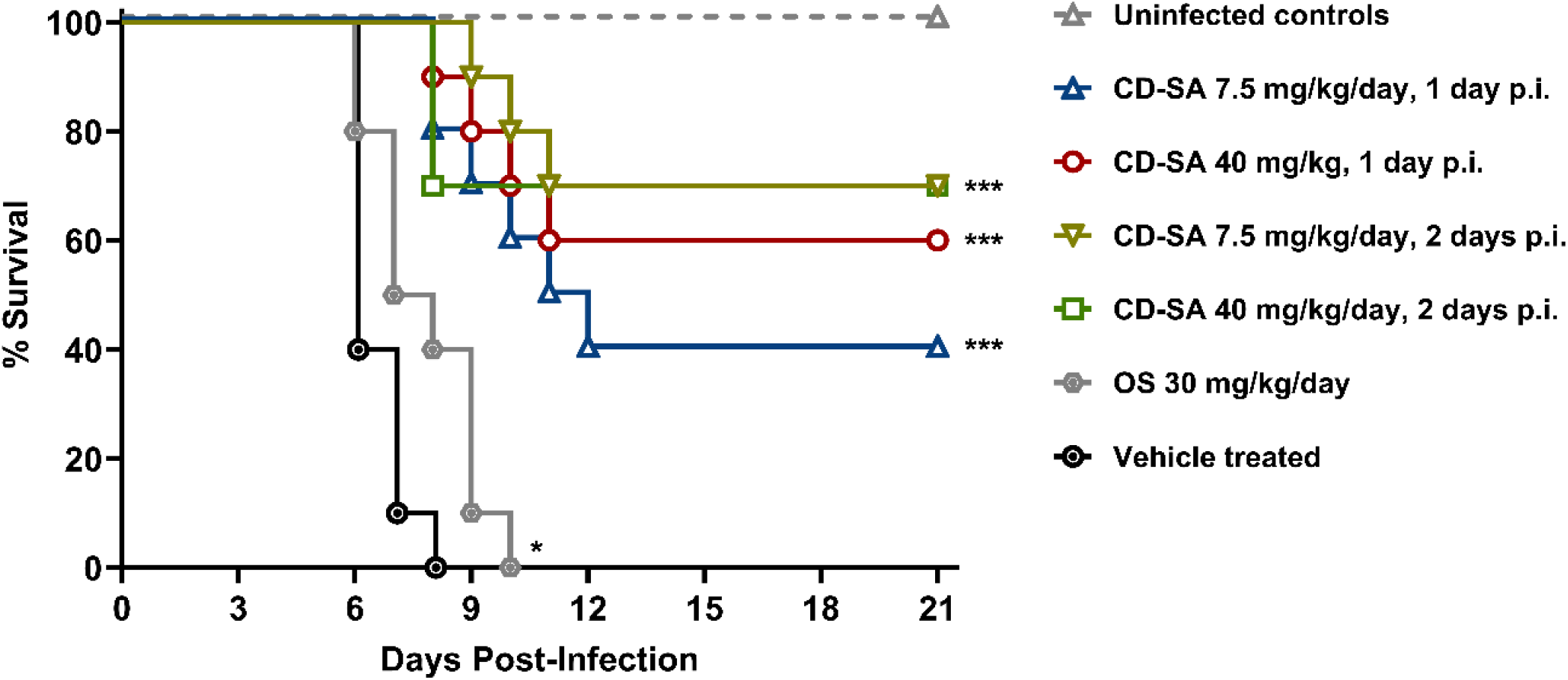
Efficacy of CD-SA and oseltamivir against influenza A/California/04/2009 in BALB/c mice. Female mice (10/group) were infected and intranasally treated with sterile saline (vehicle) or CD-SA after 1 or 2 dpi and continued for 5 days. Control mice were treated by oral gavage with oseltamivir (OS), 2 h before infection and twice daily afterwards during 5 days. Survival curves were compared by the Mantel-Cox log-rank test using Prism 9 (GraphPad Software, San Diego, CA). ***p<0.001, *p<0.05 compared to the vehicle-treated animals.

The *in vivo* results can be explained by the fact that CD-SA destroys the viral envelope likely after interaction with the HA, as demonstrated by the by the electron micrographs (Fig.3C) and fluorescence release assay (Fig.3D) of IVs exposed to CD-SA.

Interestingly, the RNA release assay did not yield a significant RNA degradation after exposure to CD-SA, this is likely because the molecule does not break the interaction between the nucleoprotein and the RNA. Upon examining all electron micrographs of IVs in the presence of CD-SA, it is evident that the viral particles are thoroughly distorted, indicating a probable release of their contents due to the mechanism of action of CD-SA. This mechanism explains the virucidal effect shown in Fig.2, and ultimately the survival curves of mice treated with this molecule. Interestingly, CD-SA or CD-6’SLN are active against both IVA and IVB Yamagata strains. However, the action against recent H3N2 variants turns out to be only virustatic. Although IVB, IVA H1N1 and H3N2 all use α2-6 bound SA as a receptor, the branching of the latter might be more complex and variable among H3N2 strains or alternatively the envelope structure of recent H3N2 strains may be more resistant to external pressures [37]. Additional development is needed to reach broader virucidal activity against all human infecting IV strains.

OS and Baloxavir Marboxil are FDA approved drugs used to treat IV infected patients. However, mutations conferring resistance to these compounds have been documented [38-40]. Here, we observed that the emergence of resistance mutations against CD-6’SLN and CD-SA after respectively 3 and 6 passages in Calu-3 cells, indicating that the barrier to resistance is higher for CD-SA than for CD-6’SLN (Table 2). We also identified that mutations conferring resistance against CD-SA (K211N/R221K) differed from those found in the presence of CD-6’SLN (Q223R), the latter being known to cause a switch in receptor usage from α2-6 bound SA to α2-3 bound SA [41]. HA normally binds to longer oligosaccharides that terminate with SAs, with a high specificity to the sequence and connectivity between the sugar monomers. For example, avian strains of IV bind to α2-3 SA on lactosamine and mammalian strains to α2-6 linkages. In using a SA directly connected to an aliphatic chain, the evolutionary pressure to adapt to the compound was no longer naturally occurring as a connectivity adaptation, but rather to a SA presented directly to the binding pocket. This can explain the higher barrier to resistance observed for CD-SA when compared to CD-6’SLN as well as the different resistant mutations identified. This higher barrier to resistance urges for further investigation of this macromolecule as a promising antiviral agent that targets HA. In addition, as shown previously for OS [16], combining CD-SA with IFNλ1 prevented emergence of antiviral resistance over nine serial passages (Table 2). Combination therapies coupling virucidal compounds and endogenous antiviral molecules with low inflammatory profile, may thus represent promising approaches to limit emergence of IV resistant variants.

In conclusion, CD-SA shows enormous potential as a novel anti-IV antiviral. Its virucidal mechanism of action correlates with *in vivo* efficacy, and further optimization of this compound and development towards the use against influenza can introduce a valuable addition to the toolkit to manage the morbidity and prevent the mortality caused by influenza infection, thus enhancing the preparedness for a pandemic.

## 5. Acknowledgments

We acknowledge Sophie Clément Leboube for her critical reading of the manuscript.

## 5.1 Supplementary Materials

Supplementary information about experiments and materials can be found in the Supplementary file.

## 5.2 Author Contributions

**Arnaud Charles-Antoine Zwygart**, Conceptualization, Methodology, Investigation, Writing - Original Draft, Writing - Review & Editing, Visualization.

**Chiara Medaglia**, Conceptualization, Methodology, Investigation, Writing - Original Draft, Writing - Review & Editing.

**Yong Zhu**, Methodology, Investigation, Writing - Original Draft, - Review & Editing, Visualization.

**E Bart Tarbet**, Investigation, Review & Editing.

**Westover Jonna**, Investigation, Review & Editing.

**Clément Fage**, Review & Editing.

**Didier Le Roy**, Investigation

**Thierry Roger**, Resources, Review & Editing.

**Samuel Constant**, Resources, Review & Editing.

**Song Huang**, Methodology, Resources, Review & Editing.

**Francesco Stellacci**, Conceptualization, Resources, Review & Editing.

**Paulo Jacob Silva**, Conceptualization, Writing - Original Draft, Writing - Review & Editing, Visualization.

**Caroline Tapparel**, Conceptualization, Resources, Writing - Review & Editing, Visualization, Supervision, Project administration, Funding acquisition.

## 5.4 Data Availability Statement

Data will be made available on request.

