## Supplementary materials for "Development of Broad-spectrum β-cyclodextrins-Based Nanomaterials Against Influenza Viruses"

### 1. Method

#### 1.1 CD-SA Synthesis

##### 1.1.1 Synthesis of β-CD-(-S-C11-COOH)$_7$

All reagents were dried before use under ~2.5x10$^{-2}$mbar at room temperature for 48 h; dry solvents were purchased and used as received*.* Potassium tert-butoxide (tBUOK, 82.47 mmol: 8.7g, Sigma) and 12-mercaptododecanoic acid (35.28mmol: 8.2g, Sigma) were dissolved in 100mL of dry DMF under an argon atmosphere and the mixture was stirred at room temperature using a mechanical stirrer, 300rpm. Heptakis-(6-deoxy-6-iodo)-β-Cyclodextrin (4.2mmol: 8g, Arachem) was added to the mixture after one hour, the mixture was placed in an oil bath at 70°C and the reaction proceeded overnight (12 h). The crude, was precipitated into 1L of diethyl ether ($Et_2O$) and vacuum filtered with a fritted disk funnel (POR 3). The solid on the filter was collected (25g), dissolved in 190mL of distilled water. Then 380mL of ACN was added to the aqueous solution creating an off-white homogeneous suspension. This mixture was vacuum filtered (POR4) – the crude was allowed to percolate through the filter by gravity: once a flow of solvent was observed by gravity alone, the vacuum assisted filtration was turned on and the filtration went smoothly. ~12grams of an off-white solid was collected and dried. It was again dissolved in 200mL of pure water and precipitated into 0.1M HCl, to obtain the protonated species as a white powder, collected via centrifugation in 45mL Falcon tubes, followed by decantation. $^{1}$H NMR (400 MHz, TFA-*d*) δ 5.37 (d, *J* = 3.6 Hz, 1H, glucose H-1), 4.35 (t, *J* = 9.0 Hz, 1H, glucose H-3), 4.28 (t, *J* = 7.4 Hz, 1H, glucose H-5), 4.10 (dd, *J* = 9.9, 3.4 Hz, 1H, glucose H-2), 3.88 (t, *J* = 9.3 Hz, 1H, glucose H-4), 3.48 (d, *J* = 12.8 Hz, 1H, glucose H-6a), 3.24 (dd, *J* = 13.5, 7.6 Hz, 1H, glucose H-6b), 3.00–2.84 (m, 2H, S-$CH_2$-$CH_2$), 2.64 (t, *J* = 7.6 Hz, 2H, $CH_2$-COOH), 1.86 (q, *J* = 7.4 Hz, 4H, $CH_2$-$CH_2$-COOH, S-$CH_2$-$CH_2$), 1.53 (m, 14H, S-C-C-$CH_2$-$CH_2$-$CH_2$-$CH_2$-$CH_2$-$CH_2$-$CH_2$-C-C-COO). HRMS (nanochip-ESI/LTQ-Orbitrap) *m*/*z*: $[M–3H]^{3-}$ Calcd for $C_{126}H_{221}O_{42}S_7^{3-}$ 877.5133; Found 877.1148.

#### 1.1.2 Synthesis of β-CD-(-S-C11-COO-NHS)$_7$

Then, β-CD-(-S-C11-COOH)$_7$ (0.377mmol: 10g) was dissolved in 100mL DMSO under magnetic stirring until a transparent homogeneous solution formed. Then, N-hydroxysuccinimide (5.317mmol: 1.03g, Sigma Aldrich) was added followed by the addition of 1-Ethyl-3-(3-dimethylaminopropyl)carbodiimide (9.53mmol: 1.85g Sigma Aldrich), and 4-Dimethylaminopyridine (1.22mmol: 150mg Sigma Aldrich) and stirred for 12 h. The crude was washed with cold acidic water (500µL of 1M HCl into 2L of MilliQ Water kept at 4°C) and centrifuged at 5500rpm at 4°C for 5 minutes inside 45mL falcon tubes. Generally, four falcon tubes were filled with 30mL of the acidic water. Then, 10mL of the crude was added into the 30mL of acidic water into per tube, forming a white precipitate. The dispersion was centrifuged at 5500rpm, the supernatant discarded, another 30mL of the acidic water added, followed by the crude, repeating the procedure until the pellet became too large for the falcon tube. The procedure was started using a new falcon tube to complete all the 100mL of crude. Once the solid was collected, additional washes with acidic washes were performed, aiming at 5 washes per pellet. The pellets were redispersed using a sonicator bath in alternation with vigorous shaking and vortexing. As the number of washes progressed, the time of centrifugation was extended to 10 or 15min as needed to sediment all the visible solid material. Then, Acetonitrile (ACS grade, Sigma Aldrich) was used to wash the white crude. Sonication and vigorous agitation with the aid of a vortex was used to completely redisperse the crude in Acetonitrile and centrifuge it down at 5500rpm and 4°C. This was repeated until a cloudy white supernatant that could not be further sedimented under these conditions was formed. This translucent (opaque) off-white supernatant was discarded and the wash was finished using Diethly Ether (ACS grade, Fisher Scientific). The crude was dispersed in $Et_2O$ using vigorous agitation, vortexing and sonication when needed. The $Et_2O$ washed were done 5 times to completely remove the acetonitrile. The final pellets were dried inside a dessicator under high vacuum for 12 h and collected as a dry powder (~12grams). The NHS coupling was monitored by DOSY NMR and $^1$H NMR.

#### 1.1.3 Synthesis of β-CD-(-S-C11-COOH)$_{7-n}$(SA)$_n$

The collected activated cyclodextrin derivative (0.036mmol: 120mg) and the N-Acetyl β-Neuraminic acid glycoconjugate (0,072mmol: 25.4mg) were dissolved in 4mL DMSO. 100µL of a triethylamine solution (140µL TEA in 1mL DMSO) was added to the DMSO solution and the mixture was stirred overnight. The next day, the product was diluted with 0.1M phosphate buffer and concentrated using amicon filters (MWCO: 3kDA). The resulting material was dialyzed in distilled water for 3 days (the dialysis water was changed twice per day) in a cellulose membrane (MWCO: 1kDA). The resulting material left in the membrane was collected at the end of 3 days and lyophilized.

### 1.2 Cell viability assay

Cell viability in cells was assessed by MTT assay. MDCK cells (20’000 cells per well) were seeded in a 96-well plate one day before the assay. A dose range of CD-6’SLN and CD-SA (from 1.58µg/ml to 200µg/ml) dissolved in serum-free DMEM or serum-free DMEM + TPCK-Trypsin (0.2µg/ml) for MDCK cells were added on cells for 24h. After the treatment, MTT reagent (Promega) was added for 3h at 37°C according to manufacturer instructions. Subsequently, the absorbance was read at 570nm. Percentages of viability were calculated by comparing the absorbance in treated and untreated wells. Each assay was performed in two independent experiments. The cytotoxic concentration 50 (CC50) was calculated by nonlinear regression analysis [log(inhibitor) vs. response – Variable slope (four parameters)] in GraphPad prism Software.

Cell viability in *ex vivo* tissues was assessed by Resazurin treatment. MucilAir tissues were treated daily on the apical surfaces with different concentrations (from 596µg/ml to 2.38mg/ml per tissue) in 30µl. Each day, 250µl of Resazurin solution concentrated at 3µg/ml in MucilAir medium was added apically for 3 h. The solution was harvested and the apical side of the tissue was washed with PBS+/+ before addition of new daily treatments. The toxicity of the treatments was assessed by comparing the fluorescence levels of resazurin (excitation at 571nm, emission at 584nm) to untreated controls.

### 1.3 Antiviral dose response assays

The viral inhibition assay was performed as previously described [1]. A dose range of molecules (from 0.5ng/ml to 100μg/ml) was pre-incubated with the virus for 1 h in serum-free DMEM at 37°C. The mix virus plus drug was then inoculated on confluent MDCK cells in a 96-well plate for 16 h. For the cell pretreatment, cells were initially exposed to CD-SA for 1 h, followed by washing and virus inoculation. In the cotreatment method, CD-SA was simultaneously added with the virus to the cells. For the post-infection condition, CD-SA was added to the cells 1 h post infection (hpi). In all settings the dose range of CD-SA was from 20ng/ml to 100μg/ml and the number of infected cells was calculated by immunocytochemistry (ICC) 16 hpi, using the primary antibody (mouse monoclonal influenza A antibody 1:100 dilution, Chemicon®) and the secondary antibody (Anti-mouse IgG, HRP-linked 1:500 dilution, Cell signaling technology). All results are presented in GraphPad Prism (GraphPad Software version 8.0, San Diego, CA, USA) as mean from two independent experiments performed in duplicate. The effective concentration (EC) 50 and 99 were calculated by nonlinear regression analysis [log(inhibitor) vs. response – Variable slope (four parameters)] in GraphPad prism Software.

### 1.4 RT-qPCR analysis, viral RNA quantification and sequencing

Viral RNA was extracted using E.Z.N.A. DNA/RNA/isolation kit (Omega Bio-tek, Norcross, GA) and quantified using RT-qPCR with the QuantiTect kit (#204443; Qiagen, Hilden, Germany) in a StepOne ABI Thermocycler as described previously[2]. For H1N1/pdm09, viral RNA copies were quantified using A/California/7/2009(H1N1) M gene *in vitro* transcripts as reference standard as previously described [3].
For the RNA exposure assay, the ∆Ct was plotted as the difference between the RNAse-treated sample and MilliQ water-treated sample.
Extracted RNAs were reverse transcribed with Superscript® III RT/Platinum® Taq Mix (InvitrogenTM), and random hexamers according to manufacturer's instructions. HA and NA were amplified with Platinum® Taq DNA polymerase (5U/ul) (InvitrogenTM)

according to manufacturer's instructions and M13-tailed primers (Supplementary Table 1). PCR purified with an MSB Spin PCRapace column (Stratec, Berlin, Germany) were sequenced through Fasteris DNA sequencing service (Geneva) and analyzed with Geneious software.

**Supplementary Table 1.** Primers and probes used for Influenza RNA quantification.

| **Primers and probes used for Influenza RNA quantification** | |
|---|---|
| **Target gene** | Influenza A M gene |
| **H1N1 probe** | FAM 5'-TTT GTG TTC ACG CTC ACC GTG CC-3' TAMRA |
| **H1N1 forward** | 5'-AAG ACC AAT CYT GTC ACC TCT GA-3' |
| **H1N1 reverse** | 5'-CAA AGC GTC TAC GCT GCA GTC C-3' |

#### 1.5 Antiviral resistance assay

H1N1/pdm09 was passaged 9 times (0.1PFU/cell) in Calu-3 cells, seeded in 6-multiwell plates, in the presence of increasing concentrations of CD-SA, CD-6'SLN, and a combined therapy of CD-SA with IFNλ or in presence of serum-free MEM only. CD-SA and CD-6'SLN were administered 1 hpi, after inoculum removal. IFNλ1 was given to the cells 24 h before infection (hbi) and again 1 hpi. After 48 h supernatants were collected, centrifuged at 3000rpm for 5min to remove dead cells and viral loads were quantified by titration in MDCK cells.

#### 1.6 Toxicity in mice

Female BALB/c mice were obtained from Charles River Laboratories (Wilmington, MA) for this investigation. The animals were quarantined for 6 days prior to use and maintained on standard rodent chow and tap water ad libitum. Mice (n=5 per group) were anesthetized by IP injection of ketamine/xylazine (50/5mg/kg) followed immediately by the i.n. mock-challenge using Minimum Essential Media (MEM). A total of 6 daily treatments of 80, 60, 20, 10, and 5mg/kg/day CD-SA or the saline placebo were administered under ketamine/xylazine by i.n. of 0.05mL beginning 1 d post-mock-infection. Individual weights of mice were recorded every day beginning on the day of mock infection and observed daily for survival through day 14. Daily weight curve graphs were made using Prism 9 (GraphPad Software, San Diego, CA).

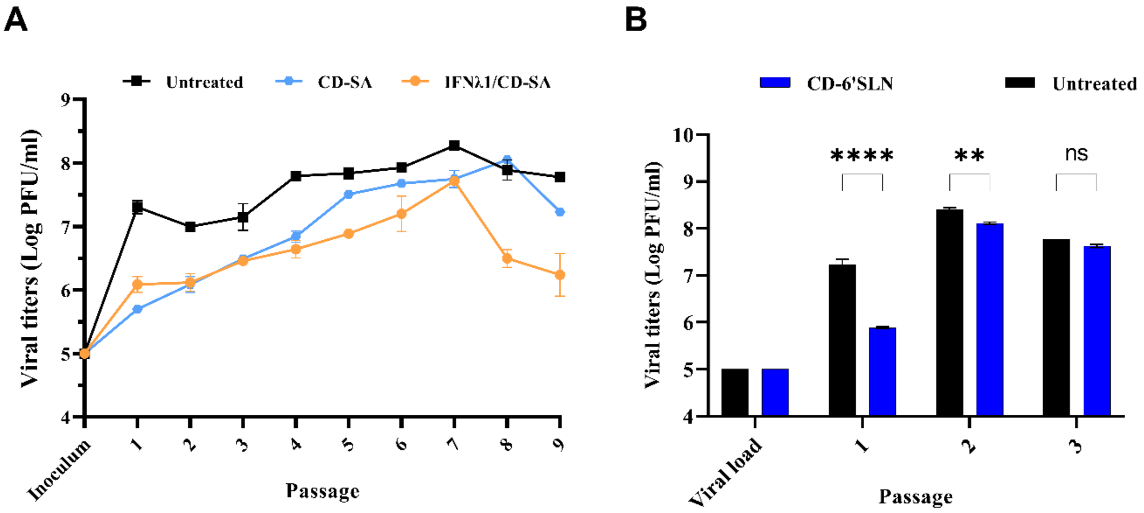


**Supplementary figure 1.** Assessment of the genetic barrier to resistance of H1N1/pdm09 against CD-SA and CD-6’SLN. (**A-B**) Comparison of viral titers from supernatants collected from Calu-3 cells infected with H1N1/pdm09 in the presence of increased concentration of CD-SA (from 1µg/ml to 173µg/ml) alone or combined to IFN-λ1 (from 15ng/ml to 960ng/ml), or CD-6’SLN (from 1µg/ml to 173µg/ml). Viral titers were assessed by plaque assay in MDCK cells and are expressed in plaque-forming units per milliliter (log PFU/ml). In **B**, statistical significance was calculated using a two-way ANOVA analysis: ****, $p \leq 0.0001$; **, $p \leq 0.01$.

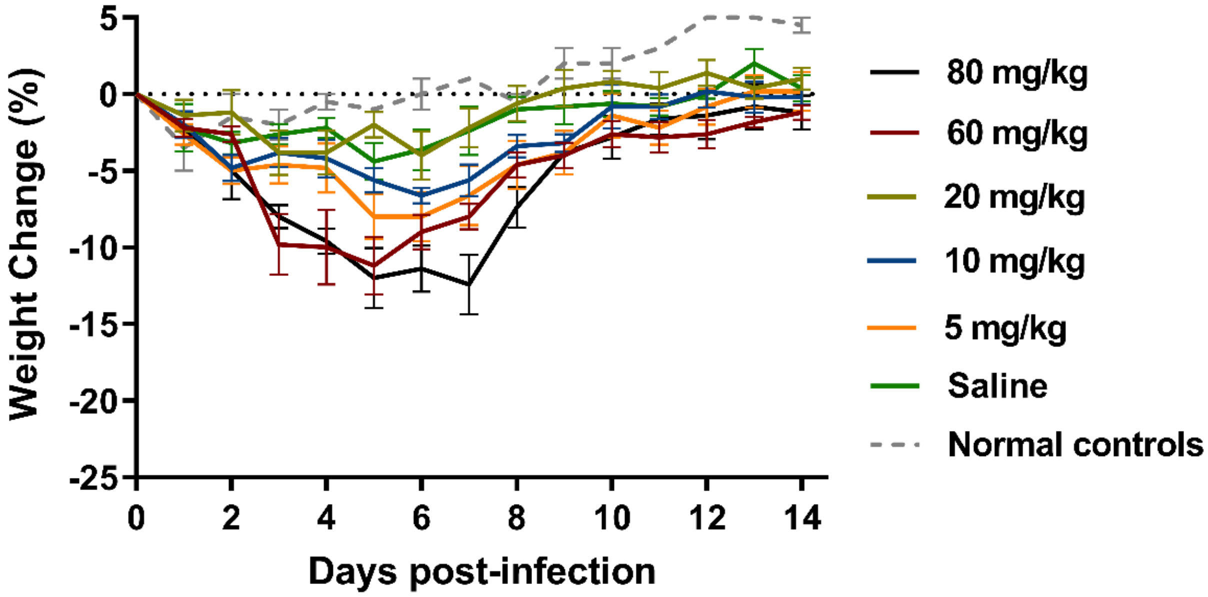


**Supplementary figure 2.** Effects of daily Intranasally treatments of CD-SA on weights of BALB/c mice mock-challenged with MEM. Weight data points represent the group mean and standard error of the percent change in weight relative to day 0. Uninfected, untreated normal controls are shown for comparison. Daily weight curve graphs were made using Prism 9 (GraphPad Software, San Diego, CA).
